# Deep Imputation for Skeleton Data (DISK) for Behavioral Science

**DOI:** 10.1101/2024.05.03.592173

**Authors:** France Rose, Monika Michaluk, Timon Blindauer, Bogna M. Ignatowska-Jankowska, Liam O’Shaughnessy, Greg J. Stephens, Talmo D. Pereira, Marylka Y. Uusisaari, Katarzyna Bozek

**Affiliations:** Institute for Biomedical Informatics, Faculty of Medicine and University Hospital Cologne, Robert-Koch Straße 21, Cologne, 50931, NRW, Germany; Center for Molecular Medicine Cologne (CMMC), Faculty of Medicine and University Hospital Cologne, Robert-Koch Straße 21, Cologne, 50931, NRW, Germany; Cologne Excellence Cluster on Cellular Stress Responses in Aging-Associated Diseases (CECAD), University of Cologne, Joseph-Stelzmann-Straße 26, Cologne, 50931, NRW, Germany; Neuronal Rhythms in Movement Unit, Okinawa Institute of Science and Technology, 1919-1 Tancha, Onna-son, 904-0495, Okinawa, Japan; Faculty of Mathematics, Informatics and Mechanics, University of Warsaw, Stefana Banacha 2, Warsaw, 02-097, Mazovia, Poland; Department of Physics and Astronomy, VU University Amsterdam, 1081HV Amsterdam, The Netherlands; Biological Physics Theory Unit, OIST Graduate University, Okinawa 904-0495, Japan; Salk Institute for Biological Studies, La Jolla, CA, USA

**Keywords:** behavior, pose estimation, kinematics, deep learning, imputation, time-series

## Abstract

Pose estimation methods and motion capture systems have opened doors to quantitative measurements of animal kinematics. However, these methods are not perfect and contain missing data. Our method, Deep Imputation for Skeleton data (DISK), leverages deep learning algorithms to learn dependencies between keypoints and their dynamics to impute missing tracking data. We developed an unsupervised training scheme, which does not rely on manual annotations, and tested several neural network architectures for the imputation task. We found that transformer outperforms other architectures including graph convolutional networks that were developed specifically for skeleton-based action recognition. We demonstrate the usability and performance of our imputation method on seven different animal skeletons including two multi-animal set-ups. With an optional estimated imputation error, DISK enables behavior scientists to assess the reliability of the imputed data. The imputed recordings allow to detect more episodes of motion, such as steps, and to obtain more statistically robust results when comparing these episodes between experimental conditions. While animal behavior experiments are expensive and complex, tracking errors make sometimes large portions of the experimental data unusable. DISK allows for filling in the missing information and for taking full advantage of the rich behavioral data. This stand-alone imputation package, freely available at https://github.com/bozeklab/DISK.git, is applicable to results of any tracking method (marker-based or markerless) and allows for any type of downstream analysis.

## 1 Introduction

Animal pose tracking was largely improved by recent progress in camera precision and synchronization, which perform accurate measurements of animal movement. On one hand, infra-red or visible spectrum cameras can be coupled with automated pose estimation (cf DeepLabCut [1], LEAP [2], DeepPoseKit [3], and their refinements for 3D or multi-animal tracking). On the other hand, motion capture systems directly register 3D positions of markers affixed on animals at key bodyparts. These pose estimation algorithms and motion capture systems estimate the position of keypoints through time, providing large quantities of data capturing animal motion. These data consist of time-series of keypoints combined into a skeleton – a simplified and widely used representation of the body for further downstream tasks such as action recognition or behavior analysis.

However both motion capture systems and pose estimation models fail to determine all keypoints at all time points in an experiment (cf Fig. 1b). In markerless techniques combined with a posteriori tracking, insufficient lightning, blurriness, animal occlusion, and automatic tracking errors result in missing or inaccurate tracked data. Some pose estimation systems, in addition to estimating each keypoint position in a given video frame, output an associated confidence score thus allowing keypoint filtering [3–8]. Marker-based techniques use motion capture cameras to directly assess the marker 3D positions and are typically more precise in determining keypoint locations [4, 9, 10]. However marker-based techniques still fail to detect all data points due to occlusions or poor quality triangulation. Independent of the cause, the missing data result in an incomplete sequence hindering the downstream behavior analysis. Since animal behavior cannot be easily scripted and additional recordings are not always possible due to constraints in experimental design, missing data is a more pressing problem in animal compared to human behavior analysis.

**Fig. 1:**
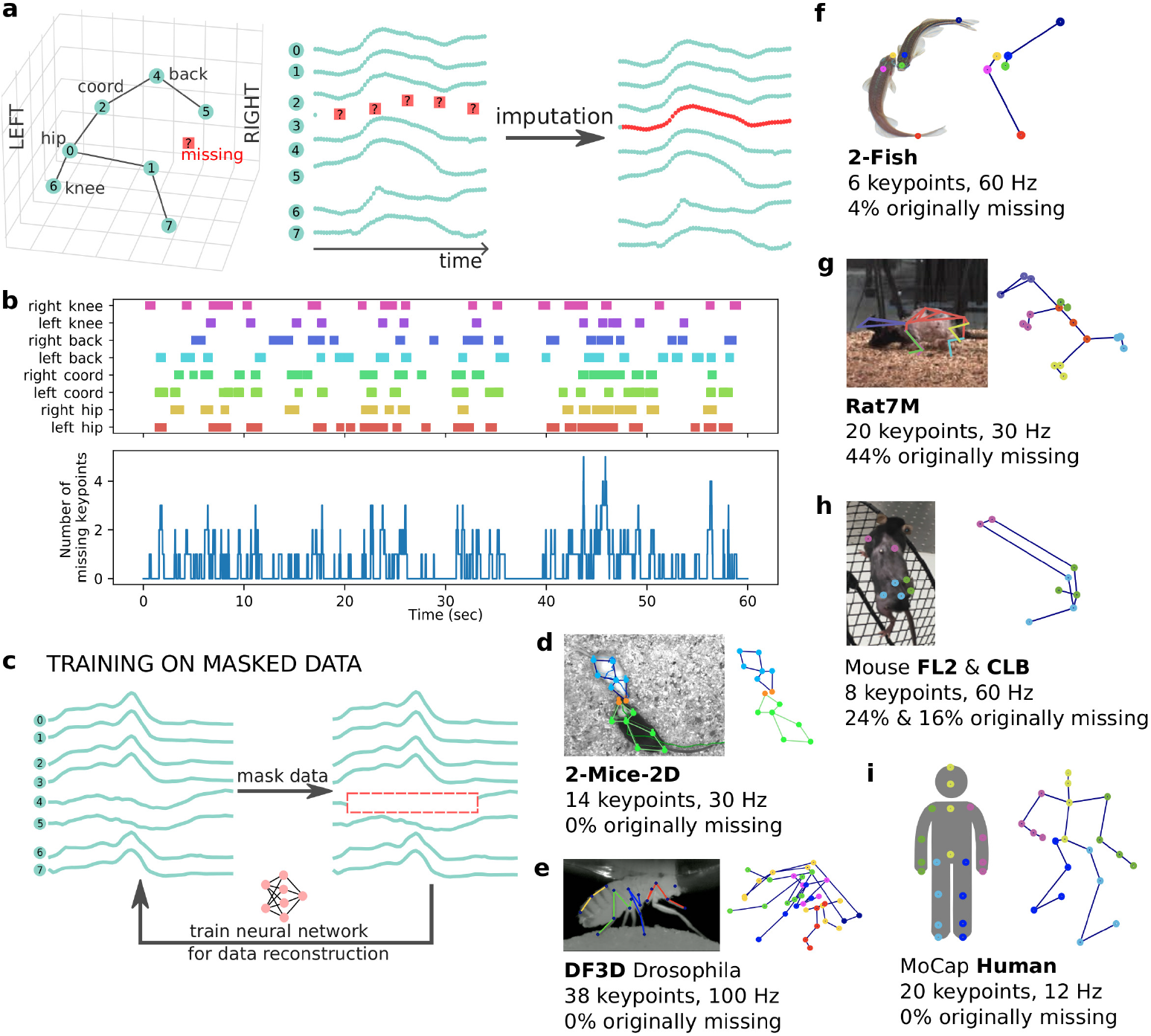
The missing data problem. a. DISK takes as input an incomplete sequence of a moving skeleton and imputes the missing coordinates. **b** Example of missing data pattern from the Mouse FL2 dataset (missing segments are indicated by colored rect- angles). During the one minute recording, the timepoints with no missing data are rare. Yet most of the time only one to two keypoints are missing, leaving the other keypoints’ coordinates as a useful source of pose information. **c** Proposed unsuper- vised learning strategy: using data segments with no missing keypoints we randomly introduce fake gaps by masking one or more keypoints and train the neural network to reconstruct the masked coordinates. **d - i** Description of the seven datasets used for the evaluation of DISK with variety of skeletons from one to two animals. The 2- Fish (**f** ), Rat7M(**g**), Mouse (FL2 and CLB, **h**) datasets show original missing data in varying proportions.

Imputing missing values in general time-series data [11–15] has been developed for applications in, e.g., medicine [16, 17], economics [18, 19], meteorology [20, 21]. Previous studies have shown that correctly imputed values can help the final forecasting or classification task [14, 15, 22]. Despite these methodological developments, none of these methods are currently used in animal behavior analysis [3–8]. Solutions for filtering of potentially inaccurate keypoint coordinates have been proposed as part of tracking packages such as Anipose and Moseq [23, 24]. These solutions are however designed specifically for post-processing of markerless tracking results. These filtering methods rely on the confidence scores of the tracking methods (cf auto-encoder in [23]) and target short-term keypoint identity swaps or jitter which represent the specific output and errors uniquely of markerless tracking methods. Importantly, **no general missing data imputation method has been developed for skeleton data** or tested on skeleton data. Skeleton data might by nature be different from other sensor data, since: (i) keypoints have physical interactions, increasing the feature correlation compared to other cases; (ii) keypoints have few degrees of freedom and many coordinates’ combinations are impossible due to physical constraints of the animal skeleton.

In this study, we propose Deep Imputation of Skeleton data (DISK), a method for imputing missing skeleton data in 2D and 3D (cf Fig. 1a). We leverage the capabilities of neural networks to use past and future information of the same keypoint, as well as the information from other non-missing keypoints, to infer missing data. We designed a fully unsupervised training scheme by introducing artificial gaps that mimic the characteristics of actual gaps in the data (cf Fig. 1c). These gaps are introduced according to the observed frequency of each keypoint being missing in the original data. This way the method is trained on gaps with similar properties as those on which inference is applied.

We show the generalizability of DISK using several skeletons with varied number of animals, skeleton representations, and behavioral tasks in 2D and 3D (cf Fig. 1d-h). The selected datasets (cf Fig. 1d-h) allow us to thoroughly test our method on different animal skeletons (mouse, fish, human, fly, and rat), varying number of keypoints (from 6 to 38), original percentages of missing data (from 0 to 44 %), and varying dataset sizes (see Table 1). Our method handles data from both markerless (2-Fish, 2- Mice-2D, DF3D datasets) and marker-based systems (Mouse FL2 and CLB, Human, Rat7M datasets), not relying on specific outputs from motion capture or tracking softwares but only on keypoint coordinates. Furthermore, DISK imputation is designed as a forecasting rather than a filtering procedure, enabling it to handle gaps longer than a few frames. Missing value imputation is crucial in animal experimentation, as recordings are scarce and repeated recordings of the same animal under the same set-up could bias the experiment as the animal has been primed. While controlled recordings of human actions can be more easily generated, we nevertheless demonstrate the capacity of DISK to perform on a dataset of human recordings as well. We show that imputed data indeed allows for detection of more behaviors and for improving the robustness of statistical comparisons.

**Table 1:** Datasets. *N kp* corresponds to the number of available keypoints, *Freq* to the considered frequency of the dataset, sometimes after downsampling (see text). The *Size train / val / test* refer to the number of 60 frames sequences (called samples) separated by a length of *Stride* in the train, validation and test sets respectively. *Missing prop* corresponds to the proportion of time points where at least one keypoint is missing.

| Dataset | N kp | Freq | Stride | Size train / val / test | Missing prop [%] |
| --- | --- | --- | --- | --- | --- |
| FL2 | 8 | 60 | 30 | 4,396 / 422 / 413 | 24 |
| CLB | 8 | 6 | 30 | 8,571 / 983 / 918 | 16 |
| DF3D | 38 | 100 | 5 | 2,095 / 652 / 614 | 0 |
| Human | 20 | 12 | 30 | 8,593 / 823 / 869 | 0 |
| Rat7M | 20 | 30 | 30 | 13,463 / 2,840 / 2,713 | 44 |
| 2-Fish | $2 \times 3$ | 60 | 120 | 99,029 / 13,327 / 15,705 | 6 |
| MABe | $2 \times 7$ | 30 | 60 | 6,820 / 986 / 622 | 0 |

Available on github^1^
, DISK can be easily trained, evaluated, and used for impu- tation on new datasets. In the field of pose estimation and motion capture methods in animal behavior research, DISK fills a gap of a posteriori imputation of missing data. Imputing keypoint trajectories in a recording will allow the scientists to take full advantage of the complex and costly animal behavior experiments.

## 2 Results

### 2.1 DISK uses an unsupervised learning paradigm

Assuming that correlations between keypoint trajectories and short-term dynamics can be learned reliably by neural networks, we designed a fully unsupervised training scheme not requiring any manual annotations (cf Fig. 1c). Several methods are designed to perform imputation on data Missing Completely At Random (MCAR) [13, 25], i.e. the value that is missing is independent of any other value, observed or not. We observed that in our datasets missing data seem to depend on the key- points, the task, and the camera set-up. For example, in the two mouse datasets, the shoulder blade markers are missing with higher probability in the 2D exploration task (FL2 dataset) while the left hip is the most probable missing keypoint in the climbing task (CLB dataset; cf Supp. Fig. A1). During training we use segments with no missing data and introduce artificial gaps that mimic the observed missing data in original dataset, in terms of gap length and identity of missing keypoints (see Methods paragraph 3.5). The gap insertion method allows the neural network to focus during training on missing data patterns similar to the ones present at inference. Such artificial gaps are created dynamically as a form of data augmentation increasing diversity of our data independent of their original size.

### 2.2 DISK imputes correctly missing skeleton data

We tested several neural network architectures – namely custom transformer (named **DISK**), Gated Recurrent Unit (GRU), Temporal Convolutional Network (TCN), Spa- tial Temporal Graph Convolutional Network (ST-GCN), and Space-Time-Separable Graph Convolutional Network (STS-GCN) – in the task of imputing a single keypoint on seven different skeleton datasets (cf Fig. 2). We additionally compared neural network solutions to linear interpolation as this method is widely used in behavior science [3–8]. Root Mean Square Error (RMSE) between the imputed coordinates and the actual coordinates within the gap, expressed in normalized units (see Methods paragraph 3.4), was used as evaluation metric (cf Fig. 2a). RMSE, calculated on the held-out test part of each dataset, is not directly comparable between datasets since the number of keypoints, their arrangement in a skeleton, and the motion range differ across animals and experiments.

**Fig. 2:**
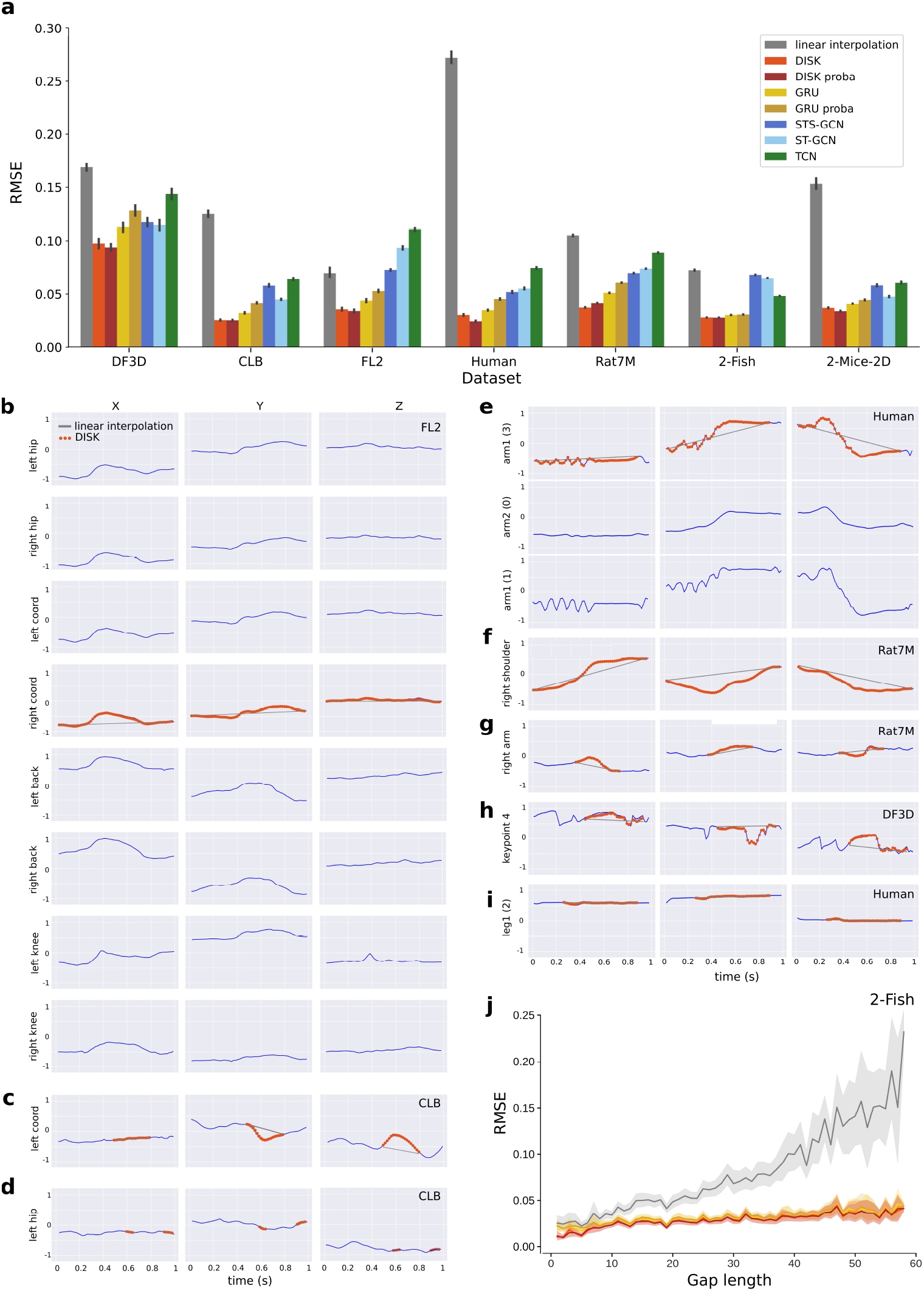
DISK and other imputation methods’ performance across datasets. **a** RMSE on all datasets for all tested architectures. ”proba” refers to a modification of the transformer and GRU architecture to additionally output a confidence interval alongside to the imputed values (see Fig. 3 and text 2.3). Fig. 2: b Example of the imputation of missing coordinates of one keypoint by DISK (Mouse FL2 dataset). **c - i** Other examples of imputed samples with different dynamics and on different datasets. Only one or a few keypoints are displayed to save space. Full plots are available as supplementary figures (cf Supp. Fig. A2, A3, A3, A4, A5, A6, and A7). **j** RMSE with respect to increasing gap length on the 2-Fish dataset by DISK, DISK-proba, GRU, GRU-proba, and linear interpolation (same color scheme as panel a).

Our method, DISK, outperforms other methods across all datasets (cf Fig. 2a), independent of their sizes and types of recorded animals (cf Table 1). Among the DL methods, GRU shows the second best performance in all except DF3D dataset, while TCN shows the worse performance in all except the 2-Fish dataset. Surprisingly ST-GCN and STS-GCN – methods developed specifically for skeleton data – do not outperform DISK. It is important to note that linear interpolation is strongly influenced by the number of static and low movement sequences in the data (cf Fig. 2i). As a result, linear interpolation performs well on the FL2 dataset that includes a large proportion of static sequences, whereas it performs poorly on Human dataset which contains only actions and no resting poses. Examples of imputation by DISK and linear interpolation are shown in Fig. 2b-i, from static poses (panels d and i) to fast movements (panels c, e, f, and h). Notably, DISK exploits correlated dynamics of specific keypoints in the imputation (cf Fig. 2b and e).

DISK displays a relatively low and constant error independent of the gap length (cf Fig. 2j for the 2-Fish dataset and Supp. Fig. A9 for the other datasets). In contrast, linear interpolation shows a major increase in RMSE and RMSE variability (cf 95 % confidence interval represented as the shaded area in Fig. 2j) with the increasing gap length, which is the reason to constrain the use of linear interpolation to small gaps [3, 5]. Remarkably, DISK does not show this trend and shows a relatively constant average error with respect to the length of the gap on all dataset (cf Supp. Fig. A9).

### 2.3 Estimated error allows to filter imputed data based on their quality

Black box predictions do not allow to assess the quality of their output. As a result, it is impossible to assess the trustworthiness of the results and thus of any downstream analysis.While crucial for real world use, there are no gold standards for uncertainty estimation of imputed data points. Existing methods [26, 27] show variable confidence intervals or low correlation with true error [28]. We adapted a recent modification of a transformer architecture [29, 30], showing good performance in estimating confidence intervals in probabilistic time-series forecasting. This modification to the transformer architecture allows the model to output, instead of a single point prediction for each timepoint and each keypoint coordinate, the two parameters of a Gaussian probability distribution over the predicted values (cf Fig. 3a and b). The standard deviation of this probability distribution allows to visualize (cf Fig. 3d and e) and assess (cf Fig. 3c) the model’s prediction confidence. The average-per-sample estimated error strongly correlates with the actual imputation error made by the model (cf Fig. 3f; Pearson correlation coefficient of 0.891 with *p − value <* 1*e −* 9, on the 2-Fish dataset – average of 5 test runs). Thus the estimated error can serve as a quality score to filter predictions based on imputation quality (cf Fig. 3c and g, and Supp. Fig. A10 for the other datasets).

**Fig. 3:**
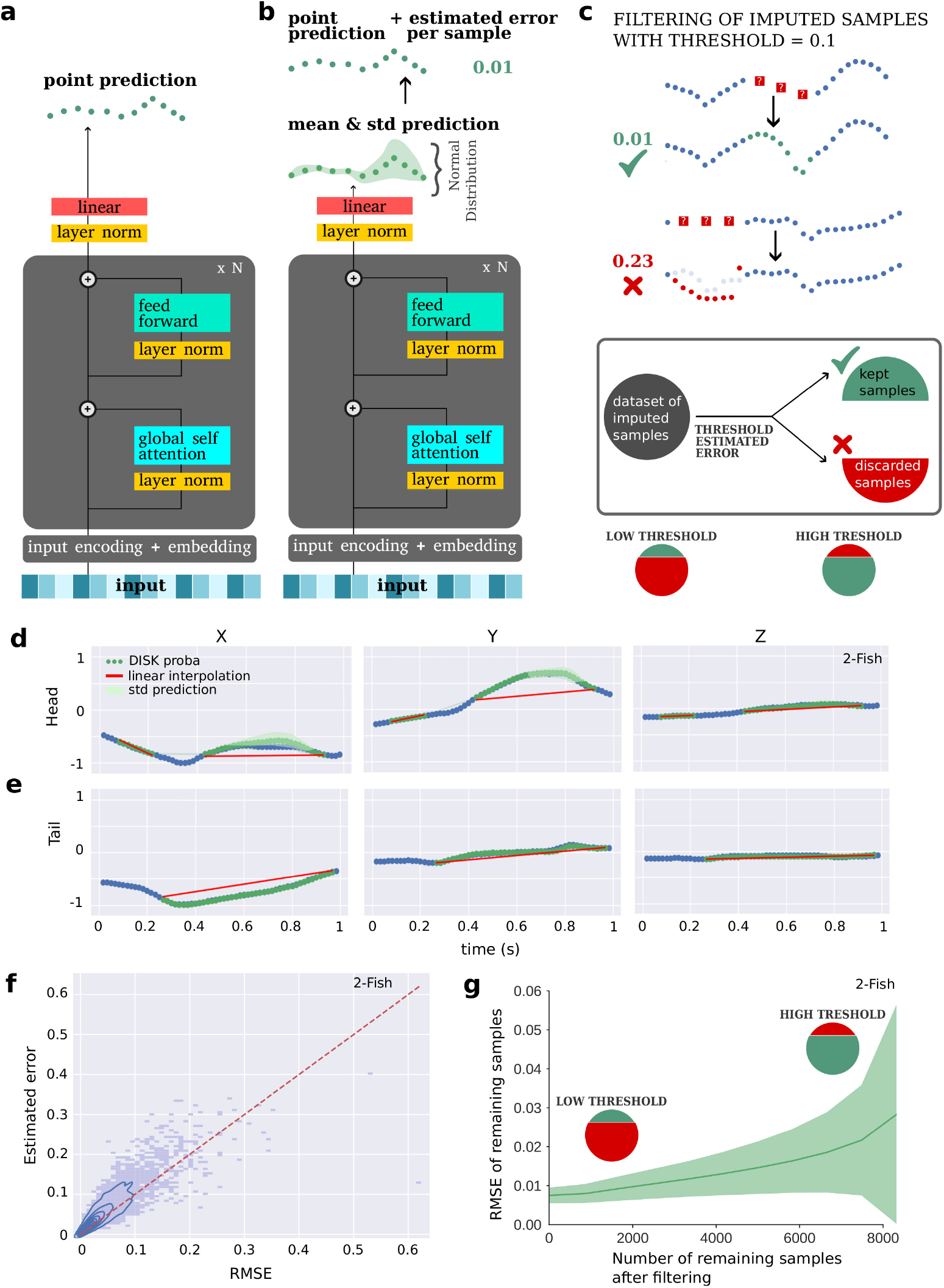
Estimated imputation error. **a** DISK architecture for keypoint coordinate imputation. This model predicts one value for each keypoint coordinate at a given timepoint. **b** DISK-proba consists of the same transformer backbone with a probabilistic head, predicting a mean value and an estimated error for each keypoint coordinate. Fig. 3:c The estimated error can be used to accept or discard samples and keep only predictions with low error for further analysis. **d-e** Examples of predictions with DISK- proba on the 2-Fish dataset. The incertitude of the prediction, shown by the confidence interval in green, increases when the prediction is less accurate. **f** The estimated error per sample correlates with the RMSE per sample on the 2-Fish dataset. **g** RMSE range of the imputed samples after thresholding the data based on their estimated error, as described in **c**. When increasing the threshold on estimated error, the number of selected samples increases as well as their RMSE.

This probabilistic component of the model can be in principle added on top of any neural network backbone. However the correlation of the estimated error with the real imputation RMSE appears much lower for GRU, the second overall best tested model. This correlation is the lowest in the CLB dataset for both GRU and DISK with a Pearson correlation coefficient of 0.43 and 0.746, respectively (cf Supp. Fig. A10).

### 2.4 DISK can impute multiple missing keypoints in single- and multi-animal set-ups

Natural skeleton poses are limited in their degrees of freedom. Observed keypoints might therefore carry important information for imputation of the missing ones. When increasing the number of simultaneously missing keypoints, the difficulty of the imputation task expands. We examined DISK in the task of imputing multiple missing keypoints and compared its performance to linear interpolation and GRU - our second best performing model (cf Fig. 2a). We trained one DISK model and one GRU model for each number of missing keypoints from one to seven on the Mouse FL2 dataset which contains eight keypoints in total (cf Fig. 4a). Both DISK and GRU show stable performance when less than half of the keypoins are missing. When more than four out of the eight keypoints are missing, the performance of both models deteriorates with every additional missing keypoint. For any number of missing keypoints DISK outperforms GRU. In contrast, linear interpolation is agnostic to the number of missing keypoints as it is applied independently to each coordinate of each keypoint and displays a constant error range.

We further analyzed the performance of DISK in the task of imputing multiple keypoints in recordings of two animals (2-Fish and 2-Mice-2D datasets). Overall, DISK shows the best performance compared to other models in this task (cf Fig. 2a and 4f) and the estimated error from the DISK-proba model correlates with real imputation error (cf Fig. 4d and e). We compared the DISK model trained on both fish to two DISK models trained separately on fish 1 and fish 2. The model trained on the motion data of both fish showed RMSE lower by 8 % to 12 % compared to the models trained on data from one fish only (see Supp. Table A2), showing that DISK is able to leverage the inter-animal dynamics to help the task of imputation (the relative improvement for the GRU model is only of 5 % to 8 %). However, since the keypoints are now positioned on two separate animal bodies, keypoint trajectories show a more complex pattern of correlation as the interaction between the two fish is expected to decrease with the growing inter-fish distance. When the two fish are close during chase or attack, they react in close loop to each other [31], but when swimming at different locations in the tank, they appear not to interact. As both fish have three keypoints and are interchangeable in this analysis, we considered different combinations of missing keypoints (cf Fig. 4b). For short gaps (length under 30; cf upper panel of Fig. 4c) RMSE stays low and increases slightly with the distance between the fish independent of the number of missing keypoints and their configurations. However, for long gaps (length above 30; cf lower panel of Fig. 4c) RMSE increases with the distance between the fish and most rapidly in cases with many missing keypoints (cases 3 and 3 + 3) compared to a single missing keypoint (case 1). This increase in RMSE with the inter-fish distance suggests that DISK strongly relies on interaction patterns between the two fish.

**Fig. 4:**
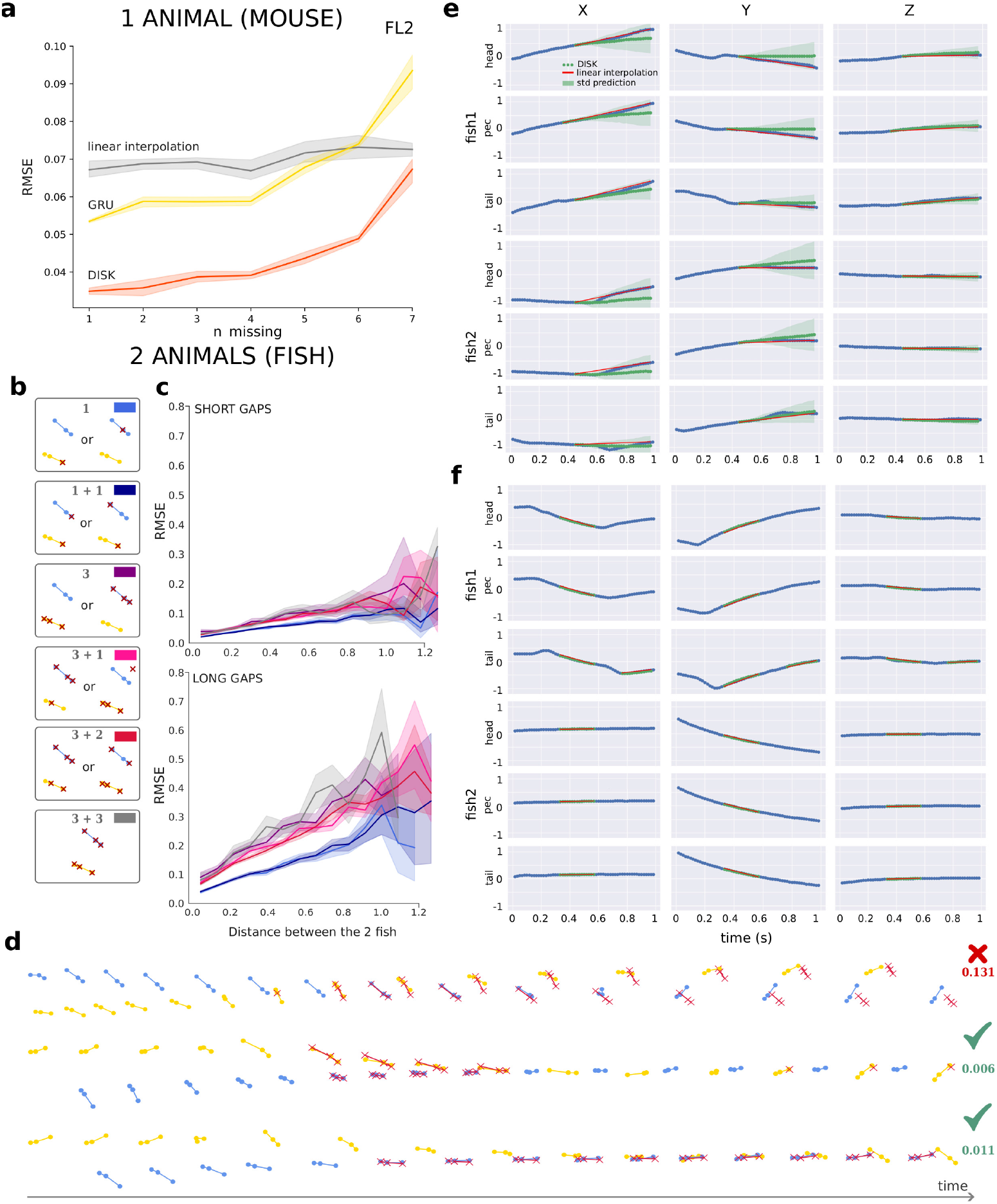
Inference of multiple simultaneously missing keypoints. **a** Imputation performance when multiple keypoints are missing simultaneously. RMSE of DISK and GRU remains low if less than 4 out of 8 keypoints in the mouse FL2 dataset are missing. We trained a separate network for each count of missing keypoints. **B - f** DISK-proba model was trained for inference of multiple keypoints in the 2-Fish dataset with uniform probabilites. For each sample dring training, a random number of keypoints was chosen as missing simultaneously, then the length of the gap was sampled from estimated observational distribution. Fig. 4:b Explanation of the legend of missing keypoints used in **c**. Both fish can have up to 3 missing keypoints. The 2 fish are interchangeable so we use the notation ”x + y” with *x ≥ y*. ”3 + 1” means one fish has 3 missing keypoints and the other one 1 missing keypoint. Color in the upper-right corner of each panel corresponds to the line colors in **c**. **c** RMSE with respect to the distance between the two fish and the number and scheme of missing keypoints for short gaps (upper panel – up to 30 frames) and long gaps (lower panel – from 30 to 60 frames). The inter-fish distances and RMSEs are expressed in normalized units as the coordinates of each behavior sequence are normalized to be in the range [*−*1, 1] as part of preprocessing for DISK (cf 3). Values in short gaps are imputed with comparable accuracy in all missing keypoint configurations, whereas in long gaps the number of missing keypoints and the distance between the two fish determine the imputation error. A smaller distance between the two fish results in a lower error, probably due to a tighter correlation between the two fish dynamics. Additionally, in long gaps RMSE is higher when all keypoints of at least one fish are missing (cases 3 and 3 + 3). We kept the cases 2, 2 + 1, 2 + 2 out to avoid clutter in the display and to compare extreme cases of few or all keypoints missing (these cases are provided in Supp. Fig. A11). **d - f** Examples of imputation of multiple simultaneously missing keypoints in a 2-animal set-up with estimated error. **d** 3D trajectories (one fish in yellow and one fish in blue) and multi-keypoint imputation (red crosses). The numbers on the right correspond to the estimated error. The first sequence corresponds to panel **e**, the second sequence to panel **f**. The last sequence shows an example with very close inter-fish distance and an almost perfect imputation. Visually the confidence interval increases with the incertitude of the network (see panel **e**). When the prediction is close to the ground truth the confidence interval is small and almost not visible on the figure (see panel **f**).

### 2.5 DISK learns coherent symmetries and similarities between motion sequences

Prediction of masked parts of the input is a core idea behind self-supervised methods [32–37]. By learning to correct the corrupted or fill in the missing content, such methods learn meaningful representations of the data. These representations can then be used for exploratory data analysis or for supervised downstream tasks such as classification. DISK might therefore not only learn to impute the missing data but also to encode keypoint sequences capturing semantic information about the underlying behavior.

To test the encoding capacities of DISK, we further inspected DISK latent representations and performed action classification on the two datasets containing labels (Human and 2-Mice-2D datasets). In the 2-Mice-2D dataset, unconstrained mice are placed in a resident-intruder scheme and three behaviors are annotated (*attack*, *investigate* and *mount* ), while remaining behaviors are labeled as *other* [38]. The Human dataset contains sequences labeled as *wash*, *walk*, *climb*, or *animal behavior*, with the majority of sequences unlabeled. Sequences from the Human and the 2-Mice-2D mouse datasets were projected and visualized in 2D with U-map (cf Fig. 5 and Fig. 6 resp.).

**Fig. 5:**
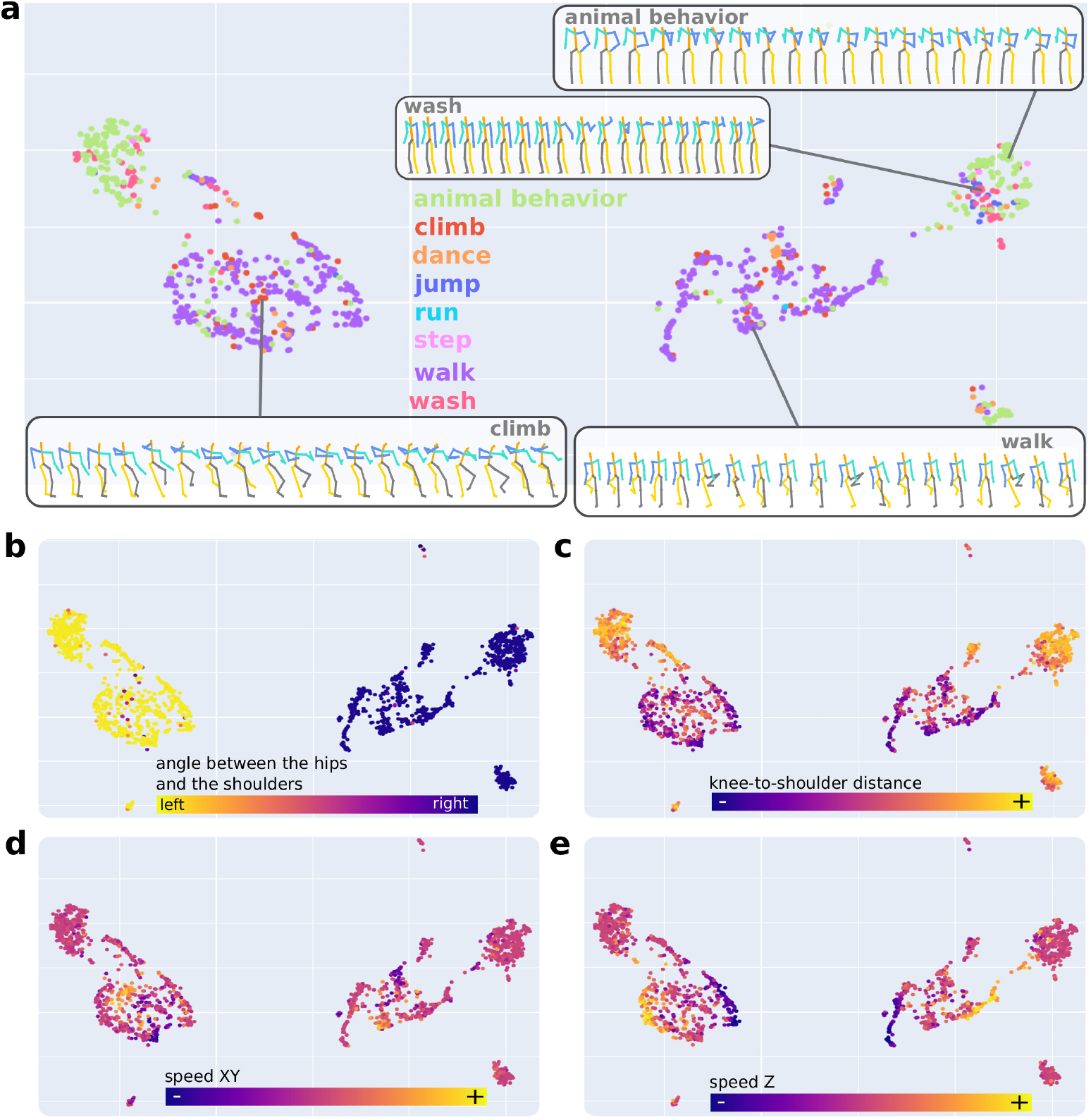
DISK learns meaningful representations of sequences from the Human dataset. a-c. Projection of DISK latent space of 10-second sequences via U- map colored by **a** the labeled action, **b** the angle between the hips and the shoulders – reflecting the global direction of the body, **c** the distance between knees and shoulders, **d** the averaged keypoint displacement in the x-y plane, and **e** the averaged keypoint displacement in the z direction. Only sequences with an action label are displayed. **a** 4 randomly selected 3D skeleton sequences illustrating 4 action types. Actions ”animal behavior” and ”wash” show similarities in arm movements, compared with ”climb” and ”walk” sequences involving more leg movement.

**Fig. 6:**
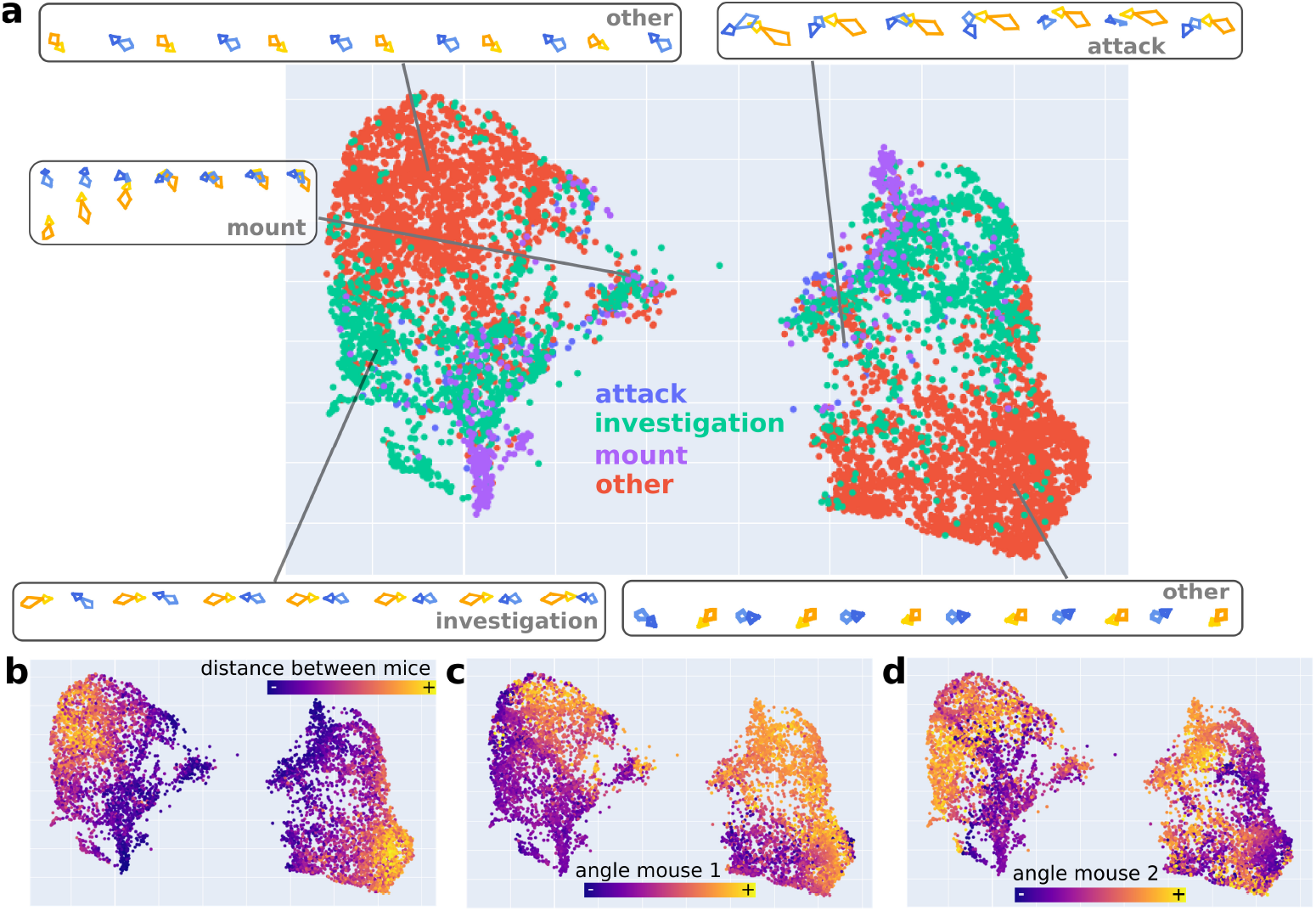
DISK learns meaningful representations of sequences from the 2- Mice-2D dataset. a - c. Projection of DISK latent space of 2-second sequences via U-map colored by **a** the behavior class of the sequence (as behavior labels are available on a frame-by-frame basis, the sequence is labeled after the majority class of the contained frames), **b** the distance between the two mice, **c** the angle of mouse 1 with respect to the vector basis, and **d** the angle of mouse 2 with respect to the vector basis. One point on the U-map corresponds to one sequence. **a** 2D skeleton representations of randomly selected sequences.

The two major groups of data points correspond to a left/right symmetry of the sequences (cf Fig. 5b). In the Human dataset the two groups of points show gradients in the distance between knees and shoulders, speed in both xy plane and in the z direction (cf Fig. 5c, d and e). In the U-map projection, sequences representing *walking* or *animal behavior* are clearly separated. Sequences representing *washing* or *animal behavior* are mixed, as well as those representing *climbing* and *walking*. These mixed sequences show similarities in arm and leg movements (see skeleton panels in Fig. 5a). In the 2-Mice-2D dataset, the two groups split sequences by the gradients of distance between the two mice and angle of each mouse compared to the vector basis (cf Fig. 6b and c). Comparing the two displayed 2D skeleton sequences from the category *other* (top left and bottom right on Fig. 6a), we can see similar interactions but rotated by 180 degrees. Most of the sequences categorized as *other* show a higher distance between the two mice compared to labeled behaviors (cf Fig. 6 a and b), suggesting their low degree of interaction. Sequences labeled *attack*, *investigation* or *mount* show markedly shorter distances between the two mice.

We further trained a Random Forest classifier for the task of behavior class prediction based on the DISK representations. In the Human dataset prediction of 8 classes showed a balanced F1-score of 0.86 and a balanced precision score of 0.94. In the 2-Mice-2D dataset prediction of 4 classes showed a balanced F1-score of 0.85 and bal- anced precision score of 0.87. These results indicate that indeed, through unsupervised imputation task, DISK has learned not only fundamental differences between motion sequences like symmetries but also higher level information characterizing different behaviors.

### 2.6 Imputed data enable finer analysis of drug-induced differences in locomotion

We next tested how can imputation of missing data improve downstream analysis of behavior experiments. We used a subset of the FL2 dataset including recordings of 10 mice performing the floor exploration task under 3 pharmacological conditions (drugs A, B and vehicle). As in the original analysis of this dataset [9], we detected steps on both sides (cf definition on Fig. 7a) and performed this detection before and after imputation. In the floor exploration task, step detection needs to differentiate between real locomotory steps and other leg movements. We therefore used kinematic rather than positional features of ankles as in [9] to detect steps. A fully-resolved step is defined as a velocity peak bordered with acceleration and deceleration events, reflecting take-off and touch-down (cf Fig. 7a).

**Fig. 7:**
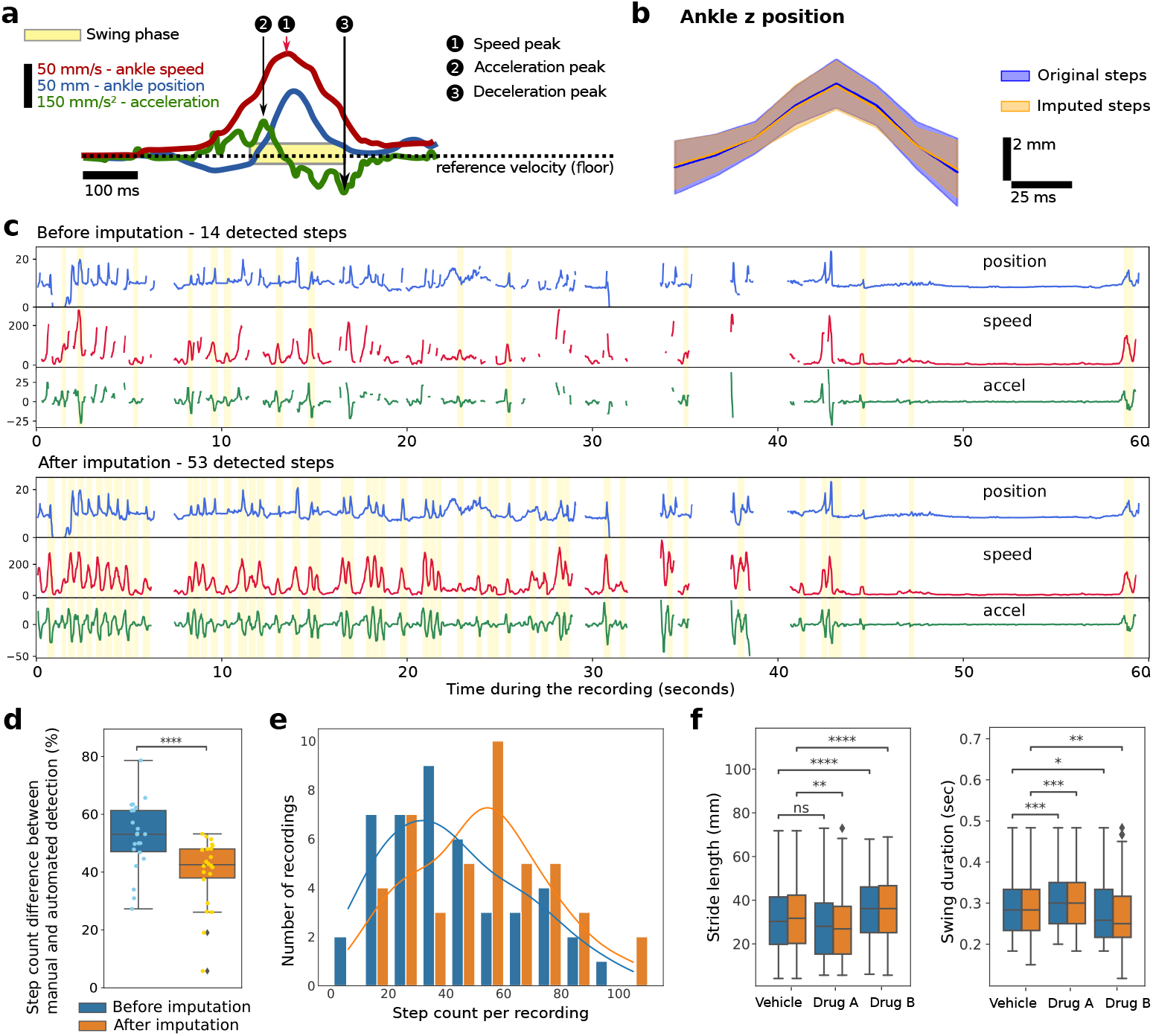
DISK allows to detect more steps and emphasizes differences in step kinematics between different treatments. **a** Step detection principle. A swing phase of one ankle is detected as a peak on the ankle speed (1). The beginning and end of the sing phase are then defined as a peak on the acceleration before the speed peak (2), and as a valley in the acceleration after the speed peak (3) (cf Methods paragraph 3.8). **b** Distribution of the step ankle trajectory for original and imputed steps. The solid line represents the mean, the shaded area covers one standard deviation above and below the mean. In orange are the steps detected on imputed data, excluding the steps corresponding to the ones detected in the original data. **c** Example of a recording with detected steps before / after imputation. More steps (53 versus 14) are detected after imputation. Remaining gaps after imputation can be due to segments with all keypoints simultaneously missing, a too long gap or an imputation quality below the selected threshold. **d** Difference between manual step counts and automated step count before / after imputation in percent. The lower the better: after imputation we miss 40 % steps on average compared to manual annotations, while before 53 % steps were missed with up to 79 % missed steps in some files (paired t-test, p-value = 7.7*e^−^*^5^). Fig. 7: **e** Steps per 1-minute recording were counted. The histogram displays the number of recordings falling into each bin of type [x, x+10) steps. After imputation there is no recording with less than 10 detected steps, and a global shift of the distribution (see solid lines) towards more detected steps with a mode shifting from 30 to 60 steps per recording after imputation. **f** Distribution of two features of the step swing phase: stride length (total distance traveled by the ankle during the swing phase) and duration. The stars correspond to the p-value of the corresponding independent t-tests. We see an increase of difference between vehicle and drug A in the stride length, and between vehicle and drug B in the swing duration. T-tests performed on each feature comparing before and after imputation inside the same treatment are non-significant. ns: *p <*= 1.00*e* + 00; *: 1.00*e −* 02 *< p <*= 5.00*e −* 02; **: 1.00*e −* 03 *< p <*= 1.00*e −* 02; ***: 1.00*e −* 04 *< p <*= 1.00*e −* 03; ****: *p <*= 1.00*e −* 04.

After imputation we detected 57 % more steps in this experiment on average and the number of detected steps after imputation is closer to manual counts (expert manual counts gathered in 22 1-minute recordings, cf Fig. 7e) The steps detected on the imputed segments are similar in shape than the original ones (cf Fig 7b) underlining the quality of imputation. In the particularly challenging example on Fig. 7c, with many gaps in keypoints, visibly more steps are detected after imputation. Gaps remaining after imputation are due to too many missing keypoints occurring simultaneously, prohibitively long gap, or an imputation below the chosen quality threshold. At the end of the recording, from second 45, we can see a period of immobility with no ankle movement where no steps detected.

We next performed analysis of step features and compared it between the imputad and non-imputed data. In this analysis we used recordings with at least 10 detected steps on each side. Without imputation, 2 recordings were discarded (Fig. 7e), whereas imputation allowed us to include all recordings in the analysis and to have an overall larger number of detected steps across the recordings. In Fig. 7f we display the dis- tribution of two features of the swing phase of detected steps. We found a significant difference in the swing stride length between vehicle and drug A after imputation (p- value = 0.0016), where the comparison was non-significant before imputation (p-value = 0.0566). For the swing duration we found a higher difference between vehicle and drug B after imputation (p-value = 0.0091 after imputation versus 0.0196 before). With this example analysis, we show that imputing data with DISK allows to use a larger portion of the recorded data in the analysis, and to analyse fine motions such as steps.

## 3 Methods

### 3.1 Neural network architectures

We selected several neural network backbones for testing: transformer, Gated Recurrent Unit (GRU), Temporal Convolutional Network (TCN), Spatial Temporal Graph Convolutional Network (ST-GCN), and Space-Time-Separable Graph Convolutional Network (STS-GCN). Two of them, GRU and TCN were designed for time-series analysis, but have not been tested specifically on skeleton data. GCNs have been developed specifically to handle skeleton data in a different task to ours, namely action-recognition. Transformers are currently state-of-art models in sequential data-related tasks and show highest capacity to learn complex and long term patterns in sequences [33, 39, 40].

#### 3.1.1 Gated Recurrent Unit (GRU)

We devised a bi-directional GRU-based architecture for the task of missing data imputation in time-series pose data. The model comprises *n* GRU layers followed by an output linear layer. We tested several hidden layer sizes from 32 to 512, one to three recurrent layers, and two values of dropout, 0 and 0.2, applied before the output linear layer. GRU with 3 layers, hidden size of 512, and no dropout showed the best performance. Our GRU architecture follows a previously published model BRITS [12] with two modifications: (i) we replaced the long-short term memory (LSTM) layers by GRU layers; (ii) we added an output linear layer. Replacing LSTM layers with GRU was motivated by the observation that GRUs perform equally well or better than LSTMs on small datasets [41, 42]. Adding a linear output layer did not affect the performance of GRU but allowed us to adapt the size of the network to different datasets and numbers of keypoints by decoupling the hidden and the output size. We chose a bidirectional flow to leverage information from before as well as after the gap.

#### 3.1.2 Temporal Convolutional Neural Network (TCN)

We tested a simple version of TCN with causal 1D convolutions with respective dilation rates of 1, 2, 4, 8 for the subsequent layers as in the original paper [43]. We used four residual blocks in the TCN. A block is composed of two convolutional layers with ReLU activation and 0.2 dropout. All the hidden layers have 256 units.

#### 3.1.3 Graph Convolutional Neural Network (GCN)

GCNs received much attention in the context of handling skeleton data for action recognition [44–46]. These networks use graph representation of the relationships between input keypoints. We tested two GCN architectures for the task of keypoint imputation: ST-GCN and STS-GCN [44, 45]. We parametrized ST-GCN to five layers with hidden size of the first layer 64, which decreased by a factor of 2 at each following layer. We chose a smaller hidden size for this network compared to other tested networks since this architecture does not scale well with increasing hidden size (cf Supp. Table A1). Indeed, with a larger hidden size, the network did not fit in one GPU V100 core memory which is impractical for the deployment of the method. STS-GCN was used with two graph encoding layers and two temporal convolutional decoder layers, with a hidden size of 256 and a dropout of 0.2. ST-GCN takes as input the skeleton while STS-GCN builds an adjacency matrix during training.

#### 3.1.4 Transformer encoder

Transformer architectures, and mixed architectures with graph and transformer components are used in action recognition tasks [47–50]. Our architecture was inspired by [51, 52]. Instead of performing a linear projection of one pose with all the keypoints into a single vector, we *flatten* the posture and apply the same learned linear projection layer separately on each keypoint. In the first scenario masking can only be done at a pose level and does not allow to mask individual keypoints. In the second scenario after *flattening*, masking of an individual keypoint is possible and allows the network to exploit the relationships between keypoints for imputation. Parallel to the linear projection of the coordinates of each keypoint, the time information (1D vector of time points), the keypoint identity and the missing mask are each passed through an Embedding layer, which acts like a look-up table (cf pytorch Embedding module). We provide the information about the missing coordinates using a binary mask indicating which values are missing. In the case of transformers, we use the binary mask two ways: the binary mask is concatenated to the input coordinates (similar to other architectures; cf paragraph 3.5), and the binary mask is embedded via an Embedding layer. The time embedding, keypoint identity embedding, binary mask embedding, and the linear projection are summed into the input tensor, which is fed to the transformer encoder layers. Based on our experiments we found that four transformer layers with an internal dimensionality of 128, a model dimension of 128, and 8 attention heads as well as layer norm and a small batch size (*≤* 32) gave the best results across datasets. A linear layer was used to produce the final output.

### 3.2 Datasets

We tested the different neural network solutions on seven 3D kinematics datasets: two 3D mouse datasets – *FL2*, in which mice perform floor exploration and *CLB*, in which mice climb on the outisde of a mesh wheel [9] –, a 2D dataset with 2 mice (*2- Mice-2D* ), a 3D Drosophila dataset (*DF3D* ) [53], a 3D Human motion CMU motion capture dataset (*Human*) [54], a 3D rat dataset (*Rat7M* ) [4], and a 3D dataset with two Zebrafish (*2-Fish*) [31].

The 3D mouse datasets comprise recordings of 1 minute of 22 mouse individuals under pharmacological interventions affecting their behavior [9]. Each mouse is recorded freely behaving in two different set-ups: an open field exploration (FL2 dataset, 101 recordings) and a climbing vertical mesh (CLB dataset, 102 recordings). The order of the recordings (set-up and drugs) is randomized among mice. The Qualisys motion capture system outputs coordinates for 10 keypoints in 3D at a frequency is 300 Hz which we downsampled to 60 Hz.

The 2-Mice-2D dataset corresponds to the task 1 of the CalMS21 CVPR 2021 challenge [38] accessible on the AICrowd website^2^
. Keypoints trajectories from top-down views of 2 mice under a resident-intruder scheme are available. We only considered the train split from the challenge containing 70 recordings from 3.2 seconds to 12 minutes at 30 Hz. Frame-by-frame annotations of four possible behaviors (attack, investigate, mount, or other) are available.

The DF3D dataset from [53] consists of videos of tethered flies with different genetic backgrounds. Immobilized at the thorax, their limbs can move a floating ball. This set-up generates a very close and precise view of the animal with seven synchronized cameras. The tethering restricts possible behaviors to a few, such as walking forward and backward, grooming, and resting. In total, 199 videos of 9 seconds each at a recording frequency of 100 Hz are available. 38 keypoints on the legs, body and head are tracked via pose estimation algorithms on the 2D videos and their 3D positions have been reconstructed in previous work [53].

We used the subset of CMU Human MoCap dataset [55] curated by [56]^3^
. Human actors perform different actions including running, walking, jumping, climbing. This dataset contains 1,577 motion sequences of more than 60 frames with 20 3D keypoints at a frequency of 12 Hz.

The Rat7M dataset consists of the 3D motion capture recordings [4], and is available online [57]. Freely behaving rats are recorded in a transparent round arena at 30 Hz with six synchronized cameras. The seven recordings depict five different animals, and have a length between 33 minutes and 2 hours 10 minutes.

The 2-Fish dataset consists of 22 videos of 6 pairs of adult male Zebrafish recorded during 90 minutes or longer with 3 synchronized cameras (2 side views and one top-down view) at 100 Hz. Three bodypoints (head, pectoral fin, and tail) are tracked with SLEAP [2] on the top-down view before animal id tracking with idtracker.ai [58] and 3D view reconstruction. The recordings start with the introduction in the fish tank of two foreign fish. Only recordings displaying rigorous interaction (maneuvers, chase, fight) were kept by the original authors [31].

### 3.3 Data preparation

Input data are composed of 3D coordinates of body keypoints. In all the datasets except Rat7M, entire recordings were separated between the train, test, validation sets: a given recording was not partitioned among the sets but assigned entirely to one of the sets. In the Rat7M data, only seven 3D recordings from five animals were available. The lengths of the recordings were very variable, from a few minutes to several hours. Separating these animals’ recordings into train, test and validation sets would result in highly unbalanced sets. Instead, we split each sequence in 70 / 15 / 15 % of the frames. The starting 70 % were assigned to the train set, the next 15 % to the validation set, the final 15 % to the test set. With this partition, there is no overlap between sets and the train set contains only prior information to the test or validation sets.

Once assigned to sets, the recordings were split into 60 frame-long sequences with a stride depending on the dataset (note the different frequencies used for each dataset in Table 1). The stride – the shift between two consecutive sequences – was set to 30 (half a sample), except for the smallest dataset DF3D where the stride was set to 5 to obtain a larger number of sequences, for 2-Mice-2D where the stride was set to 60 and for the large 2-Fish dataset where the stride was set to 120 to decrease the number of sequences. We will refer to these 60 frame-long sequences as samples. These samples are direct inputs to the neural networks. The number of samples in each dataset is reported in Table 1.

Only complete samples without any missing values were kept for training and evaluation. We tested the approach developed in the SAITS paper [13]. All available samples – even those with missing data – are included in the training set, and additional gaps are introduced on top of the existing ones. SAITS loss consists of two parts: one part evaluating the reconstruction on non-missing coordinates, and one part evaluating the imputation on the artificial gap. Although this approach increases the number of available samples for training, it did not improve the imputation in our experiments.

### 3.4 Input sequence normalization

We applied two transformations to the input samples: a *ViewInvariant* rotation, and a normalization [59, 60]. The *ViewInvariant* transformation rotates a sequence so that, in the middle frame of the input sample, the body always faces the same direction. While not removing all variability this rotation prevents the neural network from learning the position or rotation of the body in the sample and brings its focus to the intrinsic motion dynamic. In practice the body barycenter and the dominant vector in the x-y plane in the middle frame of the sequence were used to compute the rotation aligning this dominant x-y vector with the (*x* = 1*, y* = 0) vector. This rotation was applied to all coordinates of the entire sample. We did not apply a rotation in the z axis. Subsequently a min-max normalization was applied to all the coordinates of the sequence bringing every coordinate to the -1 and 1 range.

### 3.5 Artificial gap insertion

FL2, CLB, and Rat7M datasets contain natively missing values. Different keypoints can have different probabilities of being missing, and certain gap lengths can be more probable than others. These characteristics of the missing process are dataset-dependent and we mimicked them in gap generation during training. In practice, three probability distributions are estimated: the probability of a keypoint *k* being missing 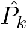for a given keypoint, the probability of a gap length *n*, 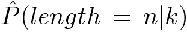, and the probability of inter-gap length 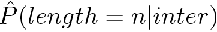. During the training, for each selected sample, an artificial gap is inserted by drawing randomly a keypoint according to the approximated 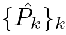, then by drawing randomly the length of the artificial gap according to 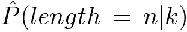, and finally by drawing randomly the starting position of the gap according to 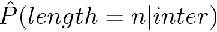.

In the DF3D, 2-Mice-2D, or Human datasets, no coordinates were indicated as missing by a specific value or as potentially missing by a confidence score. In these datasets we used a uniform distribution for 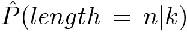. In Human and 2-Mice- 2D datasets, we also assumed a uniform probability for 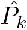. In the DF3D dataset, the 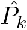 of 20 keypoints close to the fixed thorax of the fly were set to 0, as they were not displaying any position change.

We restricted the possible gap lengths to the interval [1, 58] in our samples. By capping the gap length to 58, we leave the first and last time point of the missing keypoint, allowing for comparison of imputation accuracy with linear interpolation.

Missing values were replaced with zeros - a value within the range of possible coordinate values. We therefore used an additional binary mask of the same length as the input sequence indicating with 1s where the missing values are. For all backbones except transformer (see paragraph 3.1.4) this mask is concatenated as another dimension to the input sequence, so each keypoint has 4 associated values (*x, y, z, missing*) per time frame with *missing* being 0 or 1.

### 3.6 Training procedure

We used L1 loss between the ground truth and the reconstructed gap sequence, i.e. excluding the positions and keypoints outside of the introduced gap. Each training lasted for 1,500 epochs. The network at the epoch with the lowest Root Mean Square Error (RMSE) on the validation set was saved and evaluated on the test set. All RMSEs reported in the results are calculated on the test set. We used Adam optimizer and a learning rate scheduler with rate 0.95 and steps 500. For all the networks the starting learning rate was 0.001 except for GRU it was 0.0001 as a higher learning rate showed spikes and instability in the loss during training. All the code used for this work was written in Python and Pytorch, and available at: https://github.com/ bozeklab/DISK.git.

### 3.7 Latent projection and behavior classification

For the Human and the 2-Mice-2D datasets, behavior class labels are available. For the 2-Mice-2D datasets the labels are available frame-by-frame, we then take the majority class of the frames to assign a label to a sequence. For the Human dataset, behavior class labels are given per recording, so we assign the recording label to each of its sub- sequence. For the Human dataset, we removed the samples labeled as unknown before the classification.

We used the output of the encoder layers before the final linear layer for the U- map plots. We computed the angle between hips and shoulders, the hip-to-shoulder distance, the knee-to-shoulder distance, the speed in x-y plane, distance between 2 mice, angle mouse 1 and angle mouse 2, the speed in z plane on the input coordinates without missing data. We computed the U-map projection on samples from the training set using default parameters of the *umap-learn* Python package, and projected samples from training validation and test sets.

We used a Random Forest classifier with default parameters of the *scikit-learn* Python package and fit on the latent vectors extracted the same way as for U-map. We trained the Random Forest on the same training set as DISK and reported prediction results on the test set.

### 3.8 Step detection and kinematics

We followed the definitions from [9], and used resampled keypoint trajectories at 60 Hz from the original 300 Hz recordings to do the imputation, and compute the locomotion features before and after imputation. For the imputation, a DISK-proba model was trained on multiple missing keypoints with inferred probabilities. As ankles were dropped in the FL2 dataset we pulled more data from an additional experiment to gather a sufficient data amount for the training. This additional experiment correspond to the same open floor exploration task with different pharmacological treatments and slightly different keypoints. We selected eight common keypoints: right ankle, left ankle, right knee, left knee, right hip, left hip, right back, left back. As for the FL2 dataset original data were downsampled to 60 Hz with a sample length of 60 and a stride of 30. After inspection of imputations on the held-out test dataset, we rejected imputed samples when the estimated error was greater than 0.1 for the imputation on the entire dataset.

For the step kinematics, we computed the instantaneous ankle speed (the two ankles are considered separately) and smoothed it over 0.1 second. We detected peaks on the smoothed ankle speed with a minimum height of 20, a minimum prominence of 21, and a duration in the range from 0.16 to 2 seconds. For each detected speed peak we considered potential start and end points as the closest valleys in the ankle acceleration. If the start or end points were not found, then the peak was discarded and the sequence was not considered a step. Steps whose duration were lower than 0.6 seconds, and steps where the ankle position was not lower at the start and the stop compared to the swing phase, were discarded as well. Manual counts on 22 recordings were provided by Bogna Ignatoswka-Jankowska. The swing phase features were computed as follows [9]:

- stride length: length of the 3D trajectory of the ankle or knee marker between the frames defined as swing start and swing end;
- swing duration: time difference between detected start and stop of the swing phase (in seconds);

## 4 Discussion

We demonstrate the first to our knowledge deep learning approach to impute missing coordinates in 2D and 3D skeleton data. We extensively examine performance of several neural network architectures on seven datasets of a broad range of species, including mouse, rat, Zebrafish, Drosophila, and human. The datasets differ not only in their skeleton anatomy but also in their behavioral repertoires as the Human dataset contains actions performed by actors, while the rat and the mouse behaviors are unconstrained. DISK allows to impute one or multiple simultaneously missing keypoints and shows promising results in the challenging multi-animal setting. Based on the estimated imputation error the user can filter the imputed sequences depending on their required level of tracking precision.

DISK learns spatio-temporal dependencies between keypoints and their possible dynamics. In contrast to pose estimation methods [1, 2] DISK leverages temporal information, the core component of behavior, for successful imputation. By adopting a self-supervised learning paradigm DISK does not only impute missing data but also builds meaningful representations of motion sequences. These representations differentiate sequences of varying speed, orientation and limbs motion, demonstrating the method’s capacity to learn patterns that can be used in behavior analysis. DISK is hence laying foundations for future applications of unsupervised learning to the study of behavior.

We tested DISK on two 2-animal datasets in which the animals display a high degree of interaction. In this set-up using the coordinates of both animals benefits the imputation, showing that DISK is able to learn complex, transient, and context-aware correlations between keypoints and not only correlations based on a static fixed skeleton (with for example high correlation between shoulder and elbow, and low correlation between shoulder and knee). With the fully unsupervised training scheme, further solutions for imputation of more animals separately or combined can be tested side by side and compared via the test RMSE. On the other hand DISK is not designed to impute missing data in a dataset with only one tracked keypoint per animal (e.g. centroid) such as the ones are generated by idTracker [58].

DISK relies on data alone to learn dynamics and inter-keypoint dependencies. It therefore necessitates a sufficient quantity of data for training of at least 2,000 sequences based on our experience. When creating the dataset for DISK training, the length of the samples and the stride between samples can be adjusted to influence the training set size. Longer input sequences providing more context result in better performance (cf Supp. Fig. A12c, e and f) at the cost of a longer training time (cf Supp. Fig. A12d). As alternative to the transformer backbone, GRU with the second best overall performance also shows good results on longer sequences while being faster to train and with a lower memory requirements.

DISK allows to take advantage of the entire behavioral experiment data, without the need to discard frames with incomplete skeleton information or to linearly interpolate the missing keypoints. In the Rat7M dataset 44 % of frames contain missing information, discarding them dramatically increases the per-frame cost of this type of data generation. We demonstrate that imputing data allows to recover 57 % more steps, in a subset of the mouse dataset *FL2*, leading to a finer analysis of kinematic features. Hence, recovering missing data does not only allow to access more of the data but also is fundamental to allow for comparison of precise motions such as stepping or grasping. Differences in such motions cannot be detected with general features like distance traveled or activity index.

Novel recording technologies generate more high-quality data and open opportunities to record unconstrained animal behavior in challenging, natural environments. In such set-ups, the capability of using all recorded data is particularly important, as the exact experiments cannot be repeated. DISK allows behavior scientist to fully exploit recordings of behavior whether generated in complex laboratory experiments or in unique natural conditions. The release of DISK as free and open-source Python package aims at facilitating adoption and supporting contributions and open science.

1 https://github.com/bozeklab/DISK.git

2 https://www.aicrowd.com/challenges/multi-agent-behavior-representation-modeling-measurement-and-applications/

3 problems/mabe-task-1-classical-classification/leaderboards?challenge round id=858&post-challenge=true 3accessed at https://ericguo5513.github.io/action-to-motion/#data

