## Supplementary figures and tables for "Deep Imputation for Skeleton Data (DISK) for Behavioral Science"

### Appendix A Appendix

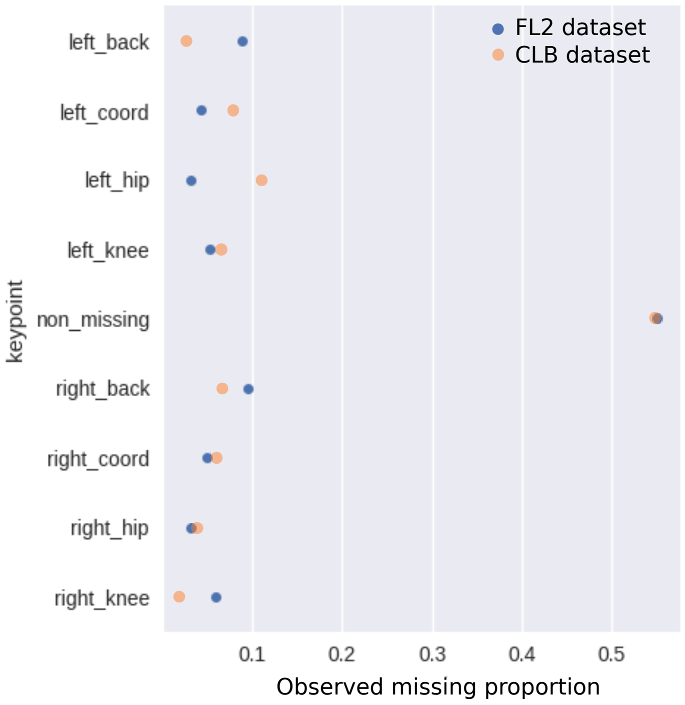


**Fig. A1**: Observed missing proportions for each keypoint in FL2 and CLB datasets. Each keypoint is considered independently of the others to calculate the missing proportion.

| Type backbone | Hyperparameters | #parameters FL2 | #parameters DF3D |
|---|---|---|---|
| DISK (transformer) | 4 layers x 128 dim x 8 heads | 408,579 | 412,419 |
| DISK-proba | 4 layers x 128 dim x 8 heads | 408,966 | 412,806 |
| GRU | 3 layers x 512 units | 11,139,096 | 11,553,906 |
| GRU-proba | 3 layers x 512 units | 11,163,720 | 11,670,870 |
| TCN | 4 layers x 256 units, kernel size=3 | 1,045,560 | 1,298,310 |
| TCN | 4 layers x 512 units, kernel size=3 | 4,055,352 | 4,523,142 |
| ST-GCN | 4 layers, 64 hidden size, kernel size=3 | 4,491,148 | 4,503,808 |
| ST-GCN | 4 layers, 128 hidden size, kernel size=3 | 17,937,452 | 17,950,112 |
| ST-GCN | 4 layers, 256 hidden size, kernel size=3 | 71,697,772 | 71,710,432 |
| STS-GCN | 4 layers x 256 units, kernel size=3 | 135,582 | 517,182 |
| STS-GCN | 4 layers x 512 units, kernel size=3 | 140,702 | 522,302 |

**Table A1**: Number of parameters for each network used in the comparison for FL2 (8 keypoints) and DF3D (38 keypoints) for an input sequence of length 60.

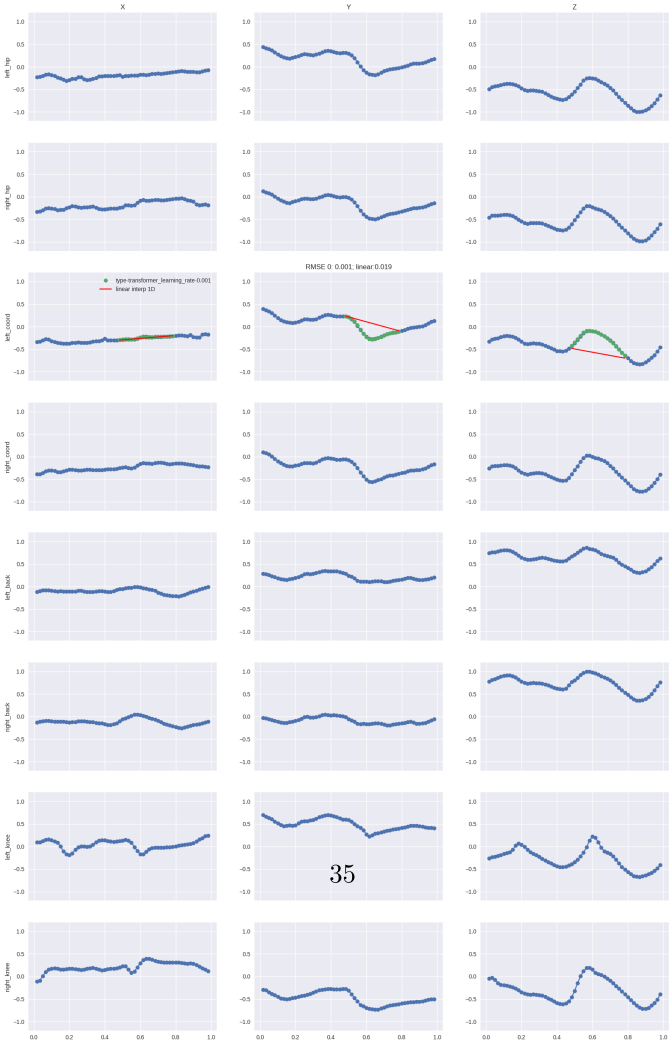


**Fig. A2**: Full plot from Fig. 2c

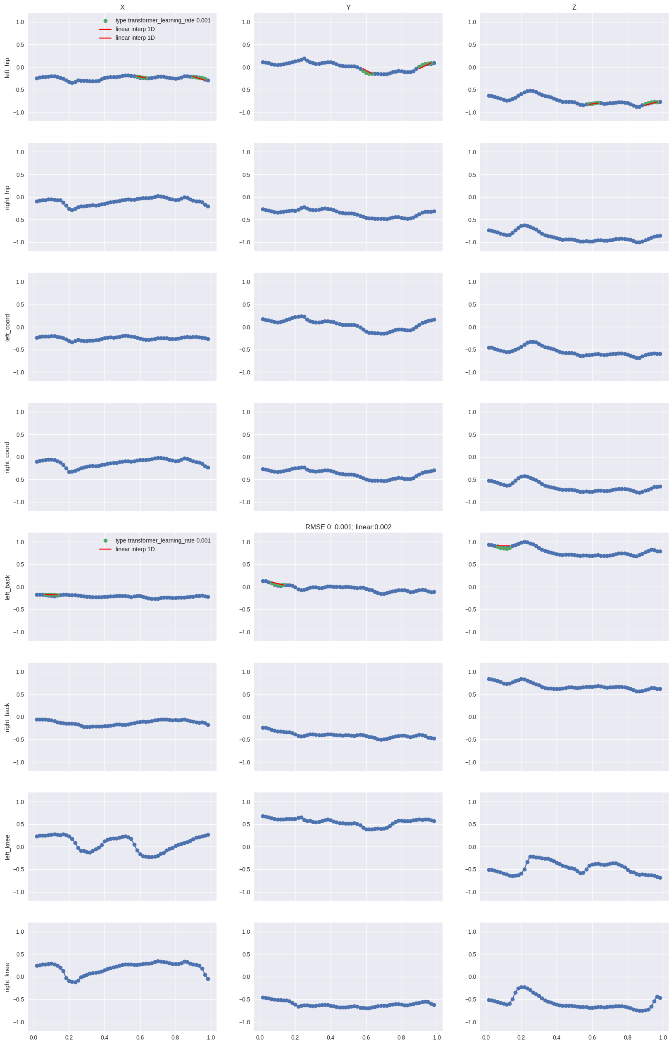


**Fig. A3**: Full plot from Fig. 2d

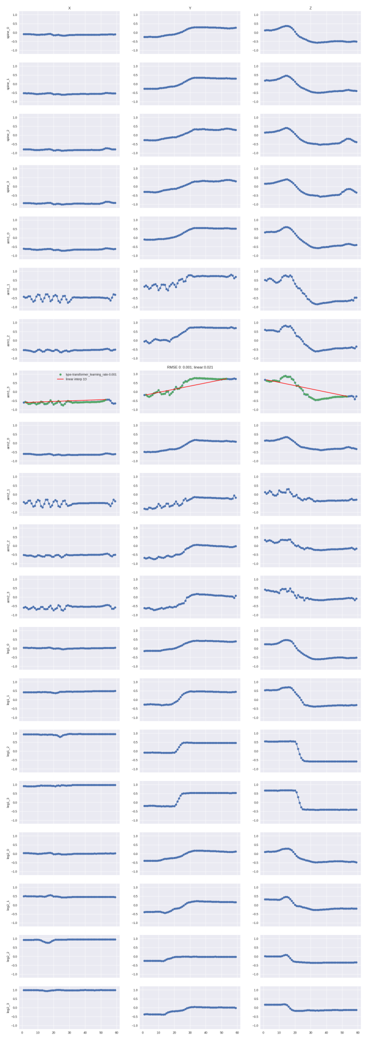

**Fig. A4**: Full plot from Fig. 2e

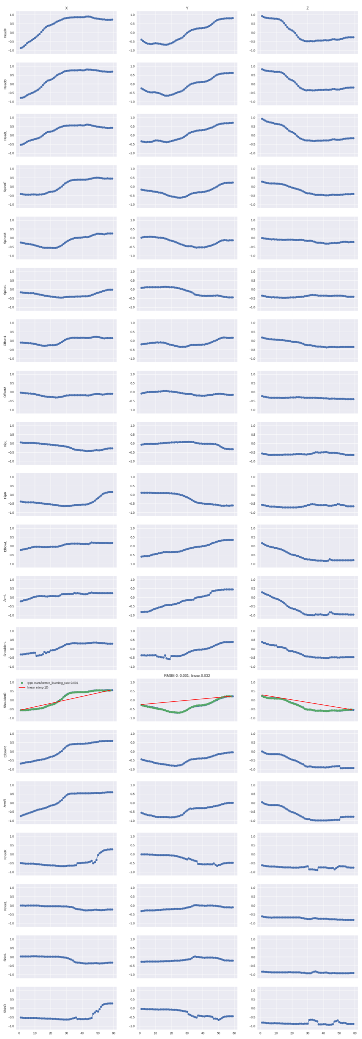

**Fig. A5**: Full plot from Fig. 2f

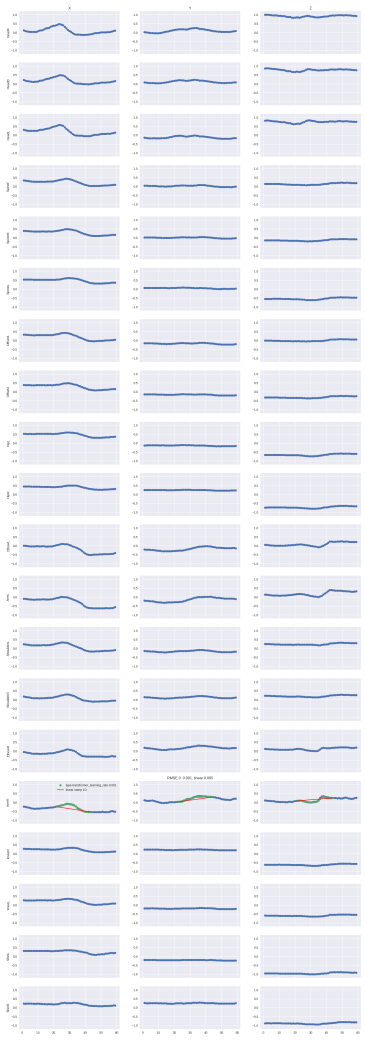

**Fig. A6**: Full plot from Fig. 2g

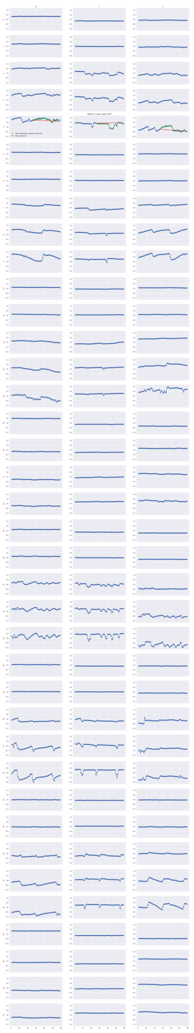

**Fig. A7**: Full plot from Fig. 2h

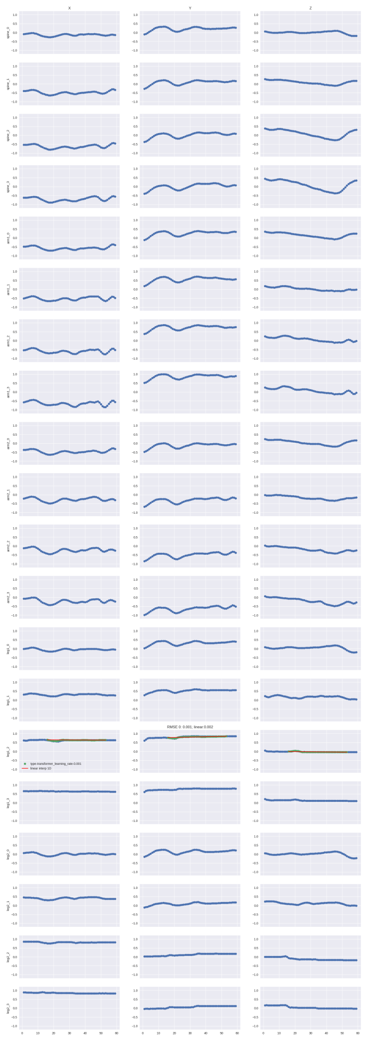


**Fig. A8**: Full plot from Fig. 2i

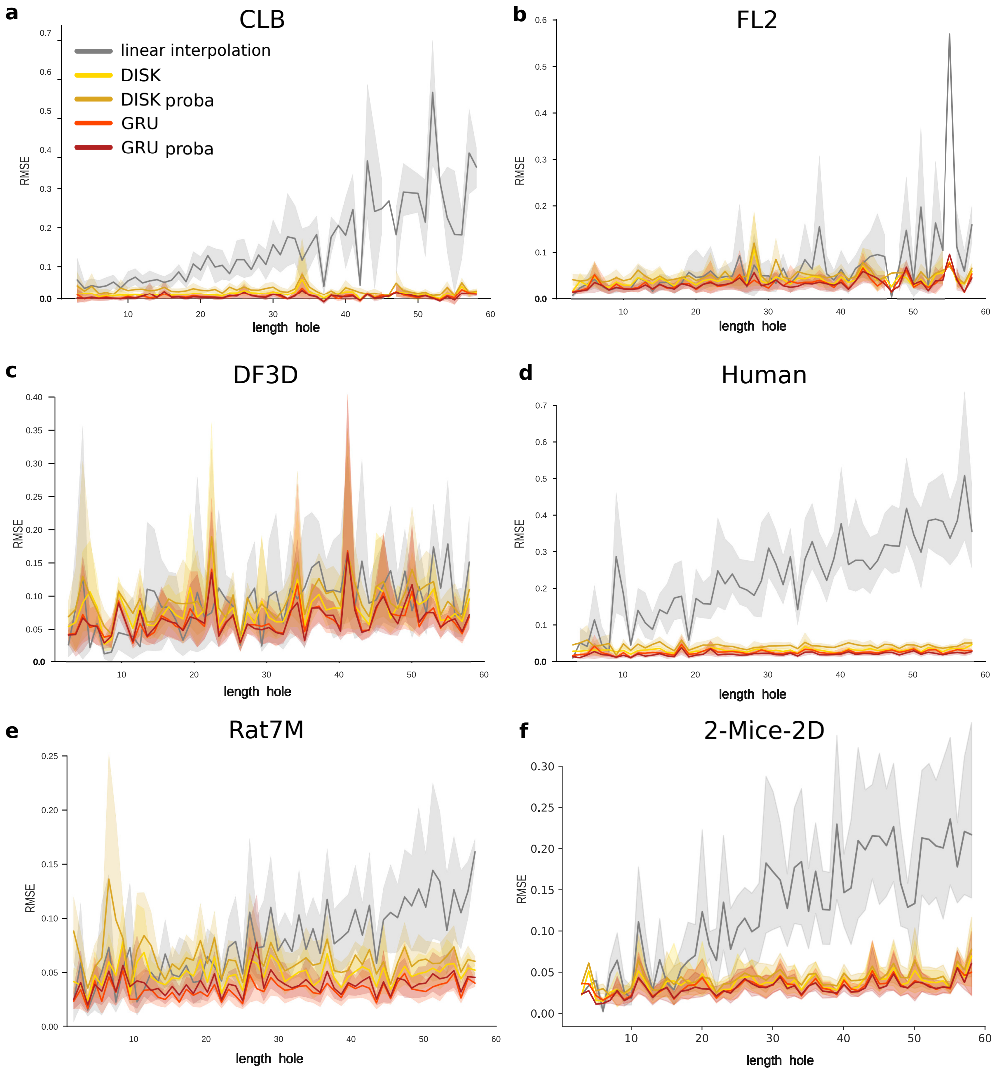


**Fig. A9**: **Test RMSE with respect to the gap length for the other datasets.** **a** CLB. **b** FL2. **c** DF3D **d** Human **e** Rat7M **f** 2-Mice-2D

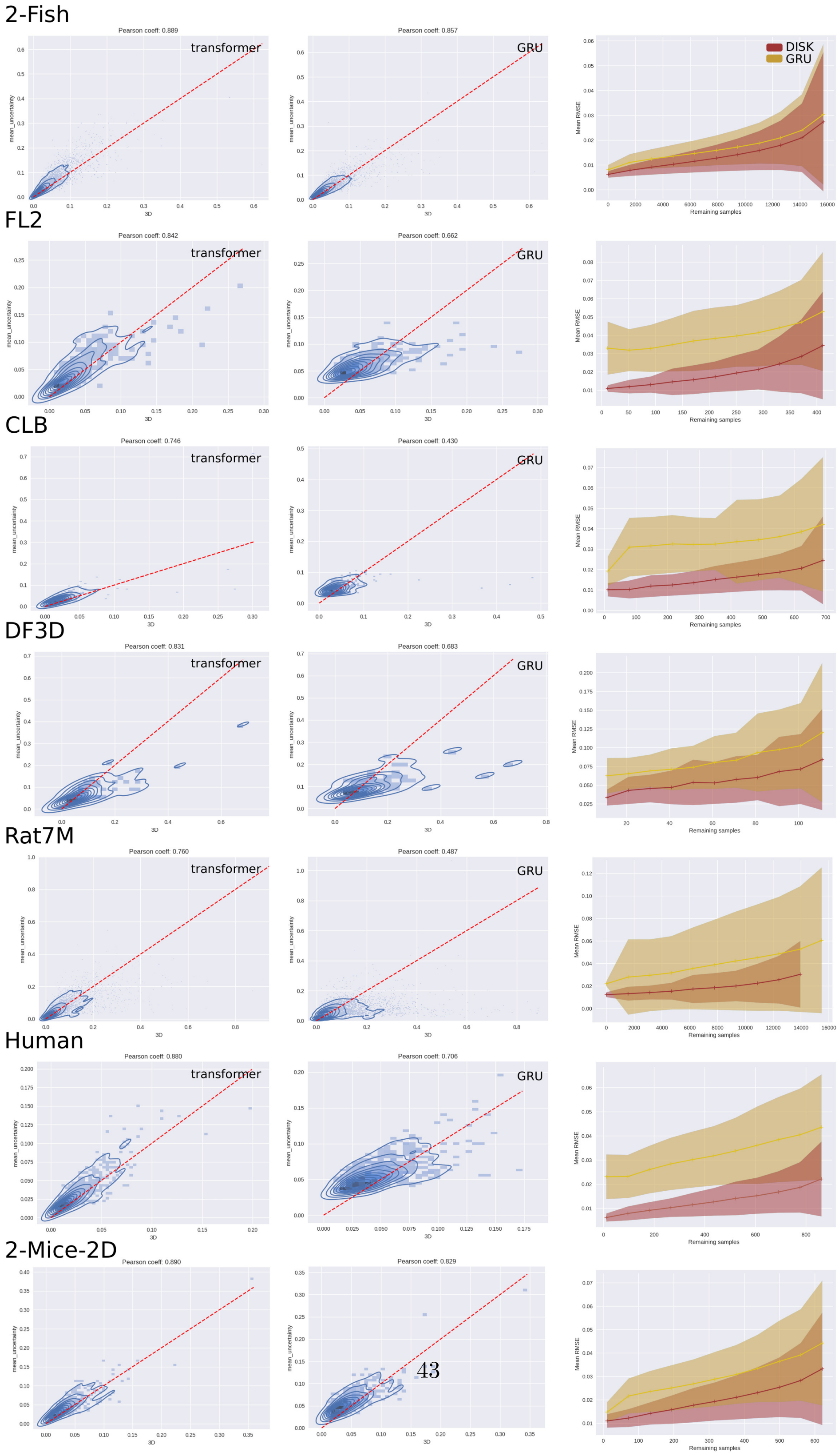


**Fig. A10**: **Estimated error correlation plots (first and second column) and RMSE after filtering based on estimated error plots (third column) for DISK-proba and GRU-proba for all tested datasets.** The red line shows $y = x$. The 2-Fish dataset gives the best correlation for both DISK-proba and GRU-proba with Pearson correlation coefficients over 0.86. GRU-proba always has a lower correlation coefficient than DISK-proba with one as low as 0.43 on the CLB dataset. The quality of the estimated error can also be seen on the third column plots where at a given remaining sample number the mean RMSE and the RMSE variance are lower for DISK for all datasets.

| Method | RMSE on fish1 | RMSE on fish2 |
|---|---|---|
| linear interpolation | 0.0826 | 0.0806 |
| GRU 2-fish | 0.0297 | 0.0319 |
| GRU 1-fish | 0.0315 | 0.0345 |
| GRU proba 2-fish | 0.0304 | 0.0323 |
| GRU proba 1-fish | 0.0323 | 0.0342 |
| DISK 2-fish | **0.0274** | **0.0295** |
| DISK 1-fish | 0.0297 | 0.0336 |
| DISK proba 2-fish | **0.0273** | **0.0294** |
| DISK proba 1-fish | 0.0294 | 0.0326 |

**Table A2**: Comparison between the models tested on the 2-fish dataset (all keypoints taken together) and the models tested separately on 1 fish or the other (one model trained on keypoints from *first* fish only, and one model trained on keypoints from the *second* fish only). Mean RMSE of 5 test runs is reported. Best models are indicated in bold.

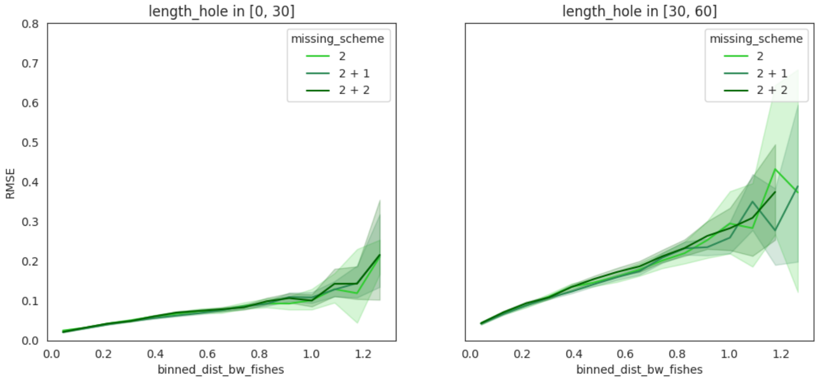


**Fig. A11**: **RMSE with respect to the distance between the two fish and the number and scheme of missing keypoints for short gaps (upper panel – up to 30 frames) and long gaps (lower panel – from 30 to 60 frames) for the cases** $2$**,** $2+1$**,** $2+2$**.** Missing cases that were left out from Fig. 4. Results obtained with a DISK model trained with uniform probability.

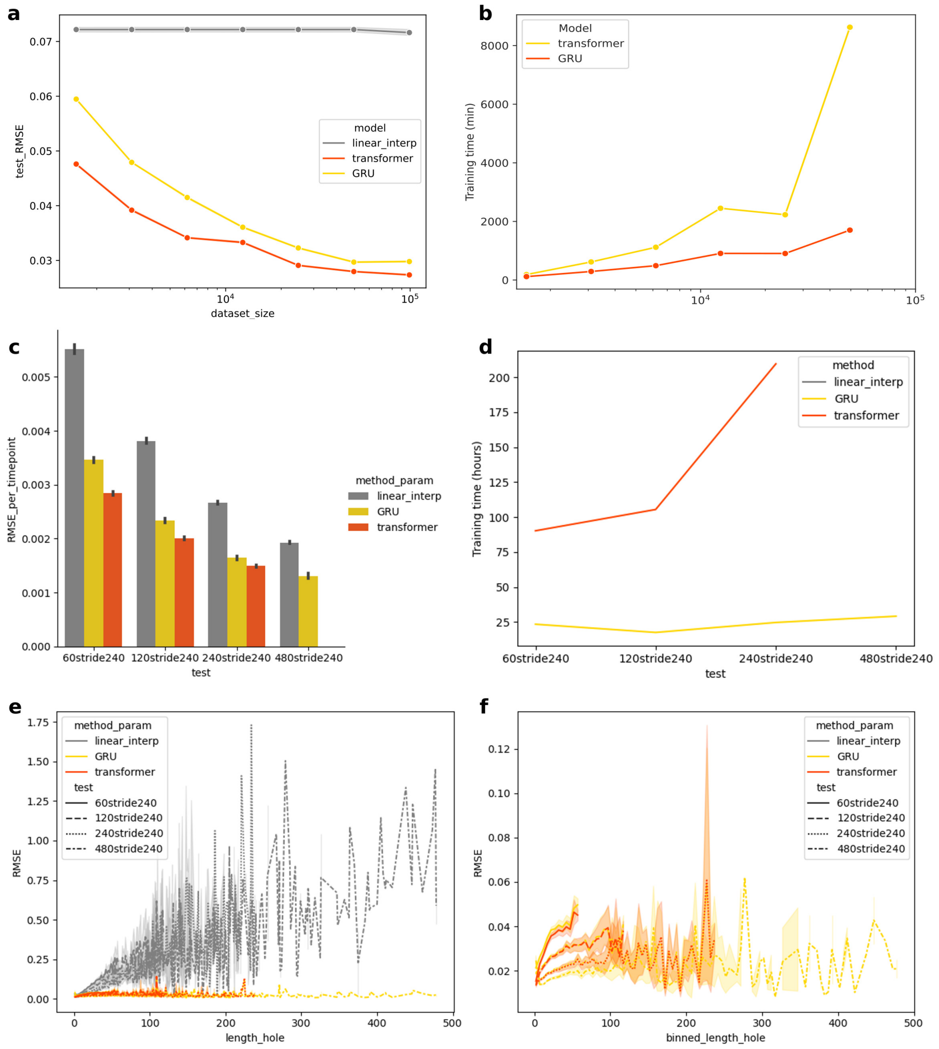


**Fig. A12**: **Further analysis on 2-Fish dataset.** **a** Performance when decreasing the size of the dataset by subsampling. The size of the dataset influences the performance: the more the better. However a division factor of 10 of the original dataset gives still satisfactory test RMSE under 0.04. **b** Training time for different size of subsampled datasets (batch size = 32) for GRU and transformer. **c** Test RMSE for GRU, transformer, and linear interpolation. An increased input sequence length while keeping the stride constant (same dataset size) show increased performance. **d** Training size for different input length (dataset size kept constant). Transformer has a big memory footprint and the model size is exceeding the memory of one GPU at 480 . On the other side, GRU's training time is kept constant for all tested input length. **e - f** Test RMSE with respect to gap length. Increased input length shows lower error, even on shorter gaps. Increased input size seems to provide a more complete picture of the dynamics and help the imputation. **f** is a zoom of **e** without the linear interpolation line.

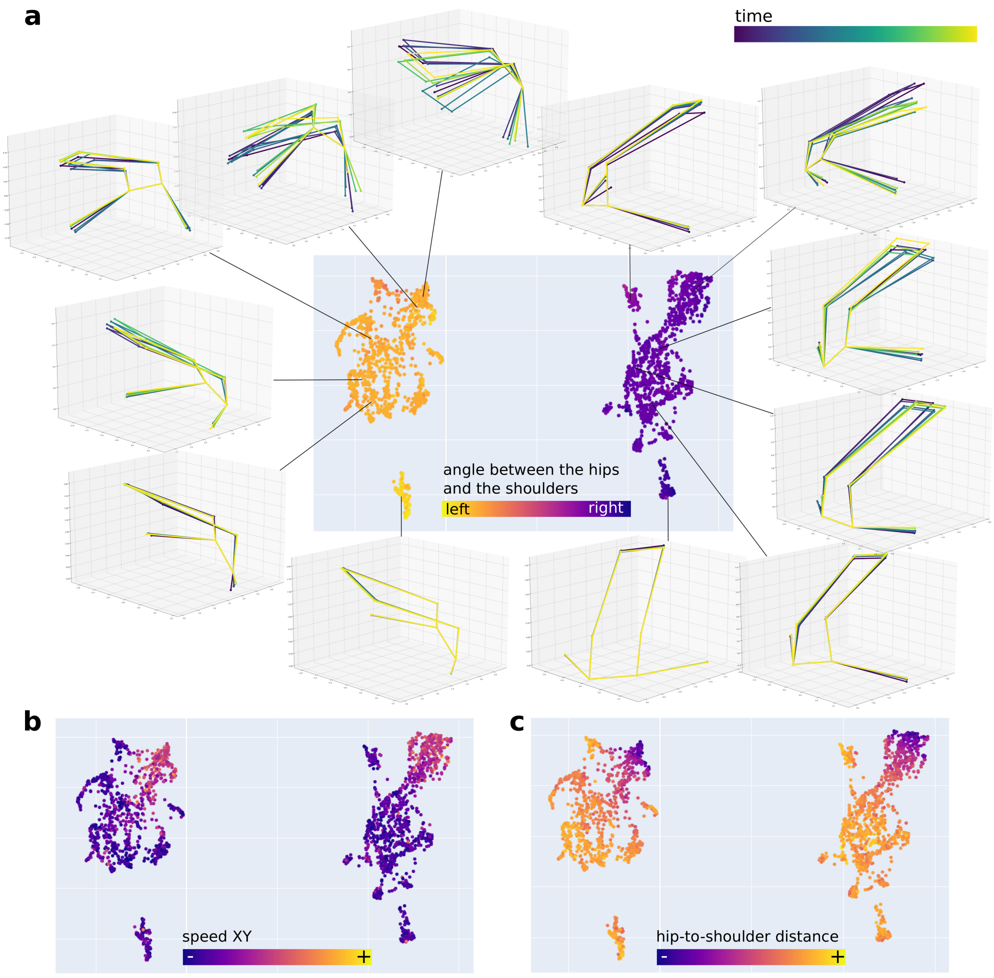


**Fig. A13**: **DISK learns meaningful representations of 1 sec-long sequences of the Mouse FL2 dataset. a - c** Projection of DISK latent space of the sequences via U-map colored by **a** the angle between the hips and the shoulders – reflecting the global direction of the body, **b** the speed in the x-y plane, and **c** the hip-to-shoulder distance – which varies depending on the posture and locomotion behavior. One point on the U-map corresponds to one sequence. **a** 3D skeleton representations of randomly selected sequences. There is gradient of increasing movement on both sides of the U-map, and clear left-right symmetry between the right and left side of the latent space.
